# Artificial intelligence model: optimizing cancer risk level predictions using machine learning and deep learning approaches

**DOI:** 10.64898/2026.08.20.745910

**Authors:** Areen Arabiat, Hamza Abu Owida, Suhaila Abuowaida, Nawaf Alshdaifat, Hamza A. Mashagba, Azlan B. Abd Aziz

**Affiliations:** Department of Communication and Computer Engineering, Al-Ahliyya Amman University, Amman, Jordan; Department of Medical Engineering, Al-Ahliyya Amman University, Amman, Jordan; Department of Data Science and Artificial Intelligence, Faculty of Prince Al-Hussein Bin Abdallah II for Information Technology, Al al-Bayt University, Mafraq, Jordan; Department of Information Technology, Faculty of Prince Al-Hussein Bin Abdullah II for Information Technology, The Hashemite University, Zarqa, Jordan; Faculty of Engineering and Technology, Multimedia University, Melaka, Malaysia; Centre for Wireless Technology (CWT), Multimedia University, Melaka, Malaysia

**Keywords:** Performance metrics, Cancer risk factor, Machine Learning (ML), cross validation, Artificial Intelligence (AI)

## Abstract

This study emphasizes the potential of computational techniques in cancer risk assessment, highlighting opportunities for specific and data-driven healthcare solutions. It examines the use of artificial intelligence (AI), machine learning (ML), and deep learning (DL) approaches to improve cancer risk assessment using a Kaggle dataset.

The study uses Java-based ML software to create and evaluate multiple predictive models, taking advantage of its powerful libraries and frameworks for processing and analyzing cancer risk indicators. This work analyzes model performance using 10-fold cross-validation, resulting in reliable generalization and accuracy estimates. Several classification techniques, such as Random Forest (RF), logistic regression (LR), decision trees (DT), Naive Bayes (NB), and Multi-layer perceptron (MLP), are used to assess their efficacy in predicting risk levels for various cancer types. To measure classification effectiveness, key performance metrics such as accuracy, precision, recall, and F1 score are produced, in addition to multi-class confusion matrices. The results show that the RF model is the best classifier for classification, with accuracy of 99.85%, F-measure of 99.80%, precision of 99.80%, and sensitivity of 99.90%. These findings demonstrate the model’s ability to effectively estimate cancer risk levels among individuals, allowing for earlier discovery and more effective medical care.

## Introduction

The continued need to determine which individuals are at the greatest risk of developing cancer (and thus most in need of screening or treatment) is evident from the high rate of cancer-related illness and death worldwide. In 2020, approximately 19.3 million new cases of cancer were reported along with 10.0 million cancer related deaths; these numbers demonstrate the size of the cancer prevention/early detection challenge that is facing healthcare delivery systems today [1]. Although the use of screening programs can significantly reduce cancer-related mortality, it generates significant downstream costs, false positives, over diagnosis, and inequitable access to and follow-up after screening for certain groups of people. The limitations and barriers associated with traditional screening approaches create a significant clinical need for a risk-based approach to cancer prevention; specifically, the identification of those at greatest risk for developing cancer and who will likely benefit from enhanced screening, chemoprevention, lifestyle interventions, or genetic evaluation and to spare those at lowest risk from additional testing and stress [1, 2].

Risk-stratification for cancer is based on the understanding that many of the factors that contribute to the development of cancer are related to potentially modifiable exposures and are therefore potentially preventable [3, 4]. Epidemiological syntheses have demonstrated that although high-penetrant hereditary mutations explain a small portion of cancers, environmental/lifestyle exposures (diet/energy balance, tobacco use, infections, occupational/exposure to environmental agents, etc.) are responsible for a large percentage of cancer burden [3, 4]. Furthermore, the concept of “risk” used in clinical practice is multidimensional, including but not limited to, absolute risk over time (e.g., screening interval), near term risk (e.g., triage), subtype-specific risk (e.g., targeted prevention), and competing risks (e.g., older age populations). Therefore, clinical decision-making tools must simultaneously consider metrics of discrimination, calibration, interpretability, feasibility of data collection, and fairness across subpopulations [5].

The use of machine learning in cancer research has provided the medical community with better tools to determine how many factors contribute to the risk of developing cancer and how these factors may interact with each other. Additionally, machine learning can help clinicians develop individualized approaches to cancer prevention and early detection by determining which risk factors most accurately relate to an individual’s likelihood of developing cancer. Traditional statistical methods do not have the same level of success when dealing with the large amount of complex data associated with the causes of cancer. Machine learning is particularly effective in analyzing large amounts of data and identifying non-linear relationships between variables [6].

In addition to its capabilities related to data analysis, one of the main advantages of machine learning is its ability to identify complex interactions between risk factors. For example, random forest (RF) and gradient boosting algorithms, among others, are able to identify linear and non-linear associations between risk factors, including specific genetic-environmental interactions that are often difficult to identify using traditional statistical models. Therefore, the application of machine learning can provide a clearer understanding of the causes of cancer [7].

For genomic researchers, the “curse of dimensionality” is a common problem that occurs when there are significantly more possible characteristics than patients available to analyze. Machine learning can be used to address this issue through the process of high-dimensional feature selection. Methods such as regression and embedded methods provide a means to quickly filter through thousands of potential biomarkers (for example, genetic variants, levels of certain metabolites, etc.) and select those that most accurately predict cancer risk [8].

Machine learning also allows for the development of accurate risk stratification and individualized predictions. By classifying individuals into different risk categories, machine learning models enable clinicians to implement targeted screening and prevention strategies. Furthermore, advanced deep learning models are able to create highly individualized risk scores based on a variety of data types including electronic health records and digital pathology images [9].

Models based on complete blood count (CBC) data have been widely used as low-friction “background” risk stratifiers that run opportunistically whenever laboratory data are being generated. Kinar et al. report discrimination performance of approximately 0.82 *±* 0.01 in two validation settings with specificity of high-80s to mid-90s depending on operating point. At a very low false positive rate (0.5%), the model yielded high odds ratios for CRC cases flagged as high-risk [10]. Thus, this line of work provides clinical rationale for passive risk enrichment using routine hematology as a proxy for subtle, pre-diagnostic physiologic perturbations (e.g., iron balance and inflammation).

From model development to real-world “flagging” workflows, a follow-on evaluation of an automated CBC-based flagging system tested operational thresholds that clinicians could reasonably act upon. In a cohort of 112,584 individuals (133 CRC cases), the top 1% risk threshold produced an odds ratio of 21.8 (95% CI 11.9–39.9) with sensitivity of 17.3% (95% CI 11.4–24.7). Meanwhile, the top 3% threshold produced odds ratio of 11.0 (95% CI 7.0–17.2) with sensitivity of 32.3% (95% CI 24.4–40.9); the corresponding “yield” (cases per flagged individuals) was 2.1 and 1.0 respectively [11]. These results illustrate a central implementation trade-off for risk factor classification: modest sensitivity at stringent alert thresholds can still be clinically valuable if the flagged subgroup is small enough to enable targeted outreach, navigation, or expedited diagnostic work-up.

A complementary CRC screening application evaluated whether CBC-based ML could enrich for colonoscopy-relevant pathology beyond invasive cancers. In a screening-colonoscopy cohort of 17,676 individuals (60 CRCs and 1014 high-risk precancerous lesions), colon flag achieved (at 95% specificity) odds ratios of 5.1 (95% CI 2.3–8.9) for CRC and 2.0 (95% CI 1.6–2.6) for advanced precancerous lesions compared to normal colonoscopy [12]. When CRC and advanced precancerous lesions were combined, sensitivity was 8.1% (95% CI 6.4–9.8) at 95% specificity and 16.8% (14.5–19.0) at 90% specificity, with subgroup breakdowns reported for average risk, family history, and personal history strata [12]. The study’s key contribution is demonstrating that the learned hematologic signature is not purely a “cancer detector,” but also enriches for clinically actionable precancerous endpoints important for prevention-oriented screening pathways.

Tabular risk modeling has also been applied to harder-to-screen malignancies such as pancreatic cancer where absolute risk is low and symptom-based presentation occurs late. An integrative study of pancreatic cancer risk prediction showed that logistic regression (LR) was able to reach an area under the curve (AUC) of 0.78 while a random forest (RF) model reached an AUC of 0.88 on training data and an AUC of 0.77 on held-out testing data. The authors used model interpretation (SHAP) to characterize feature contributions [13].

For lung cancer, where risk is strongly shaped by heterogeneous exposures (including smoking intensity, duration, cessation, and environmental co-exposures), a case-control dataset was used to compare single learners and ensemble stacking strategies built from conventional ML algorithms. In that comparative analysis, an RF achieved AUC 0.877 and an XGBoost-based stacking model achieved AUC 0.887, indicating incremental improvement from heterogeneous ensembling over a strong baseline learner [14]. This study highlights how ensemble meta-learners can improve robustness when individual base models capture different aspects of non-linear interactions among risk factors.

The application of ML to genitourinary oncology has compared it to established clinical tools that have been developed and tested. In order to gain acceptance for the use of machine learning, the comparison typically will be made to an established clinical tool (risk calculator) that has already been validated and tested, instead of “no model.”

A recent development and validation study of a cost-sensitive RF compared to PSA, PSA density, and the ERSPC risk calculator for clinically significant prostate cancer (csPCA) found that at 70% specificity for csPCA, the RF had a positive likelihood ratio of 7.8, while the PSA had a positive likelihood ratio of 6.9; the PSA density had a positive likelihood ratio of 10.3 and the ERSPC risk calculator had a positive likelihood ratio of 18.4 [15]. This example illustrates how the use of ML in risk-factor classification improves upon traditional methods by increasing the discrimination of the model and improving its ability to predict true positives and true negatives. The methodological implication for risk-factor classification of this study is that discrimination alone is insufficient, and that clinical utility is improved when the threshold for disease identification matches the acceptable miss rate for the condition being identified, and when the model translates into measurable reductions in invasive medical procedures.

In contrast to the PSA studies that quantified predictive performance through the use of receiver operating characteristic (ROC) curves and focused on the development and testing of new ML models for predicting the presence or absence of disease, studies of breast cancer have utilized tabular risk-factor classification to identify both the predictive performance of the models and the modifiable and non-modifiable determinants of the condition. A large case control study conducted in Iran (1,009 cases and 1,009 controls) examined the relationship between lifestyle and behavioral, reproductive, and sociodemographic factors and the presence of breast cancer [16].

Using three different ML algorithms (RF, bagged CART, and XGBoost), the researchers found that the area under the ROC curve was highest for the RF algorithm (AUC = 0.90) followed by the bagged CART algorithm (AUC = 0.89), and lowest for the XGBoost algorithm (AUC = 0.78); the researchers noted that several predictors including chest x-ray history, deliberate weight loss, abortion history, and postmenopausal status were important and consistent across all models. This finding aligns well with the clinical need for risk factor classification, since the models are serving as a structured analytical lens to help prioritize risk factors for targeted prevention and earlier detection strategies.

Another related study examining breast cancer incidence risk used biochemical biomarkers to develop and test multiple ML models and provide both estimates of discrimination and calibration [17]. The retrospective cohort included data from over 500 patients who underwent comprehensive evaluation for breast cancer risk, and the researchers developed and tested seven different ML model variants including LR and DT based models. They found that the best performing model variant, a biomarker based LR model, had an AUC of 0.859; they also estimated the mean AUC across all seven model variants (range = 0.779–0.862) and accuracy (range = 0.780–0.841), and provided explicit estimates of calibration and error metrics (Brier score) to emphasize the importance of evaluating the quality of the probabilities generated by the model for risk communication purposes. For risk-factor classification pipelines particularly those designed for patient facing risk stratification these calibration and error metrics are operationally relevant, since poorly calibrated probabilities can result in distorted eligibility decisions (i.e., screening invitations) even if the AUC value of the model is reasonable.

This study is going to help apply ML to classify cancer risk factors better by first identifying the clinical issues as relevant to ML, then provide concrete guidelines to implement ML in a clinical setting. The study applies a well-documented public data set, as an example, to bridge clinical objectives to implementable ML. In doing so, the study demonstrates how the most critical modeling choices need to be aligned with clinical decision-making.

The study has provided two major contributions. First, the study has provided a clinical reframing of risk factor classification as an actionable stratification task – focusing on how models can guide decisions in screening and/or prevention, rather than simply optimize predictive accuracy. Second, the study has introduced a methodological taxonomy that will distinguish between three modeling tasks; risk-tier classification, cancer-type classification, and integrated genetic-environmental stratification, and explain what success would mean for each in the clinical arena. Table 1 provides a comparison table of key aspects from previous studies.

**Table 1.** Comparison table of key aspects from previous studies.

| Ref | Cancer / endpoint | ML method(s) | Validation approach |
| --- | --- | --- | --- |
| [10] | CRC detection | CBC-based ML model (tabular) | External validation across countries |
| [11] | CRC “flagging” | ML risk score with percentile thresholds | Population-scale operational evaluation |
| [12] | CRC + advanced precancer | ColonFlag (CBC-based ML) | Screening cohort evaluation |
| [13] | Pancreatic cancer risk | Logistic regression; RF; SHAP interpretation | Train/test split (held-out testing) |
| [14] | Lung cancer risk | RF; stacking ensemble (XGBoost meta-learner) | Internal comparison |
| [15] | Prostate cancer (csPCa) | Cost-sensitive RF vs clinical comparators | External validation cohort |
| [16] | Breast cancer classification | RF; bagged CART; XGBoost | Case-control evaluation |
| [17] | Breast cancer incidence risk | LR + multiple ML variants | Internal evaluation with calibration metric |

## Methodology

The proposed model in this study evaluates the performance of multiple classifiers using Java-based ML software to examine the usefulness of approaches for predicting cancer risk variables. The data which used in this study was obtained from Kaggle [18]. The dataset consisted of 2000 records and 20 features. To work with the proposed model, the data set must be in csv format. However, the most crucial step in classifier development is pre-processing. Following the preprocessing step, the dataset is loaded into a ML model. Data classification typically depends on training and testing procedures. 10-fold cross-validation was executed to eliminate overfitting and improve the model’s performance and accuracy. RF, DT, MLP, NB, and LR classifiers were used in this study to predict cancer risk variables. Confusion matrices will be used to evaluate the findings, which include numerous metric categories such as AUC, accuracy, precision, F-measure, and recall. Fig 1 displays the system architecture for the cancer risk factor prediction model, Fig 2 depicts the proposed model for cancer risk factors using Java-based ML software while Fig 3 illustrates all attributes of the cancer risk factor prediction model.

**Fig 1.**
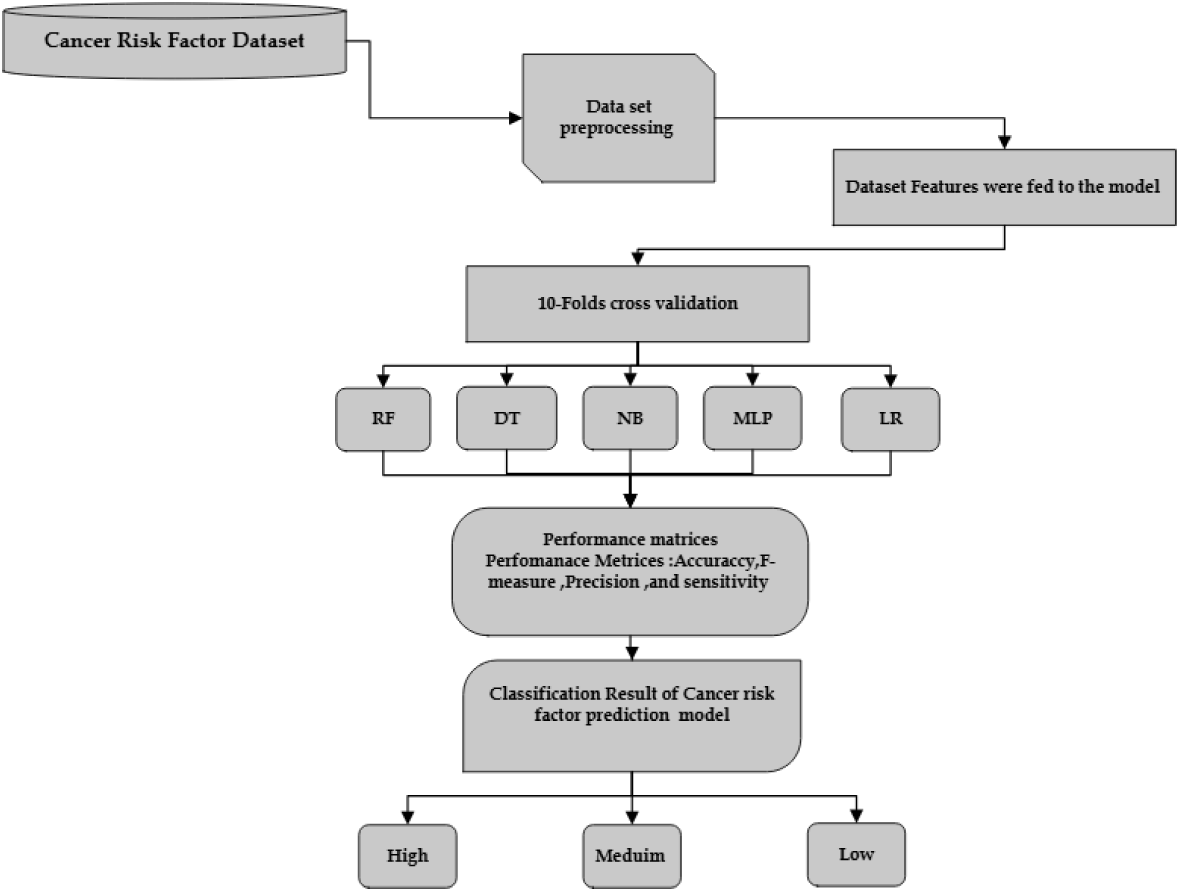
System architecture for cancer risk factor prediction model.

**Fig 2.**
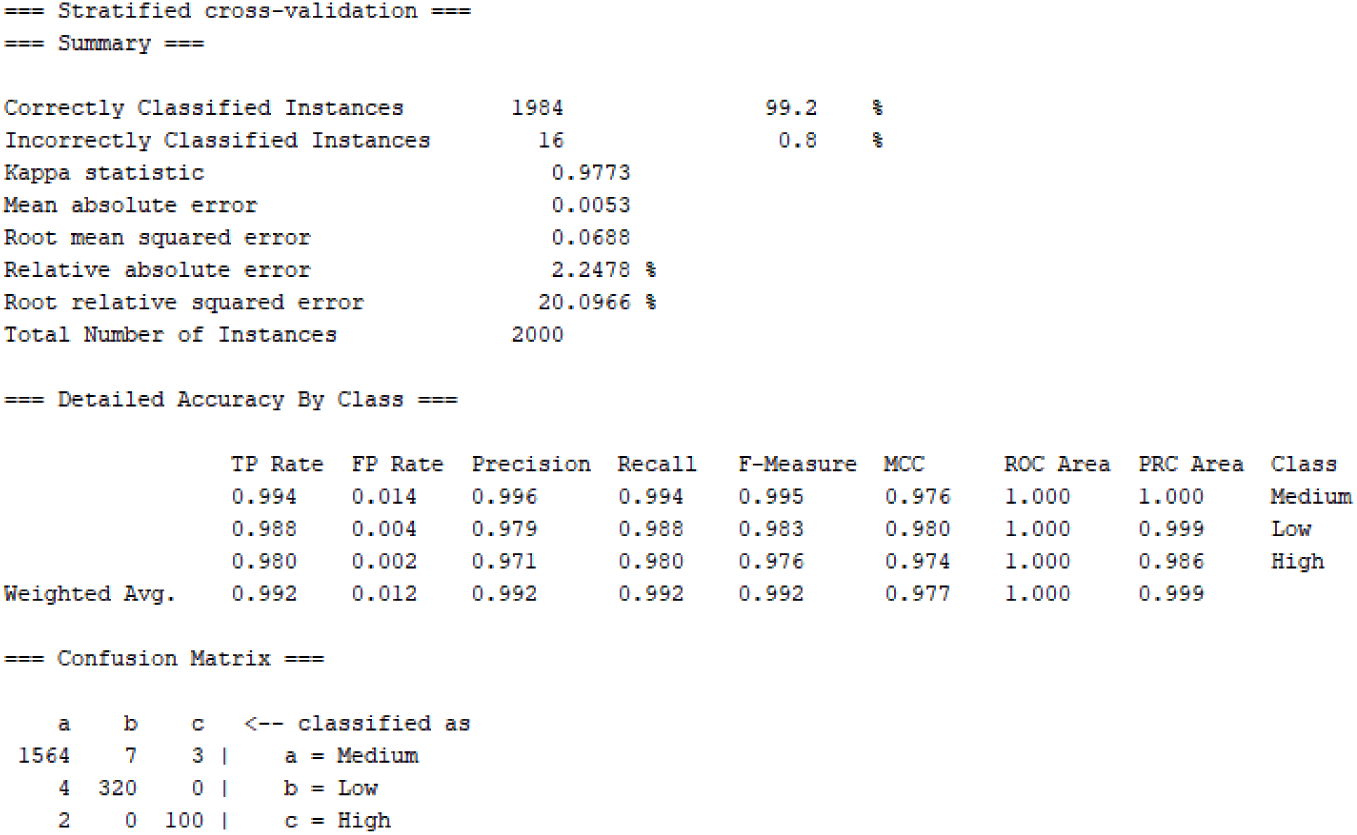
Cancer risk factor prediction model using Java-based ML.

**Fig 3.**
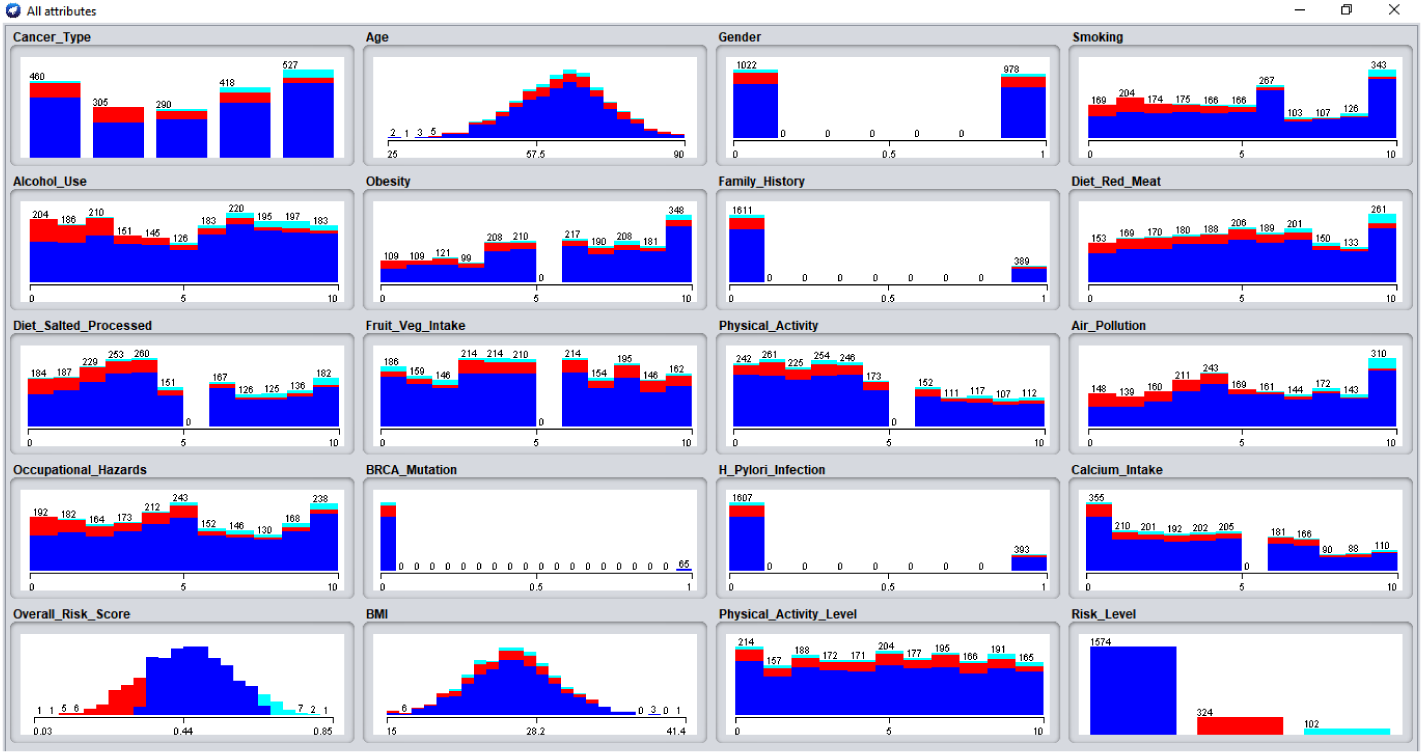
Visualized attributes for cancer risk factor prediction model.

### Dataset

This study uses a dataset of 2,000 records with 20 numerical attributes appropriate for exploratory data analysis and ML to examine lifestyle, environmental, and genetic influences on five prevalent cancer types: lung, breast, colon, prostate, and skin. The Overall Risk Score, a composite metric ranging from 0 to 1 that measures cancer risk, is a crucial component. Based on this score, the dataset divides people into three risk categories: Low (*<*0.33), Medium (0.35–0.66), and High (*>*0.66). Numerical coefficients for lifestyle and environmental factors, such as smoking habits, alcohol consumption, dietary intake, physical activity levels, exposure to air pollution, occupational hazards, and calcium intake, are included alongside demographic characteristics like age, gender (binary encoding: 0 for female, 1 for male), and Body Mass Index (BMI). Furthermore, binary indicators (0 or 1) for genetic and medical factors including H. Pylori infection status, BRCA mutations, and family history of cancer are supplied, expanding the dataset for thorough analysis. Table 2 depicts cancer risk factors attributes.

**Table 2.** Cancer risk factors attributes [18].

| Feature | Description |
| --- | --- |
| Cancer_Type | Categorical variable indicating the type of cancer (Lung, Breast, Colon, Prostate, Skin). |
| Age | The age of the individual (in years). |
| Gender | Gender of the individual (0 = Female, 1 = Male). |
| Smoking | Numeric index representing smoking habits (0–10 scale). |
| Alcohol_Use | Numeric index representing levels of alcohol consumption (0–10 scale). |
| Obesity | Numeric index representing obesity levels (0–10 scale). |
| Family_History | Binary indicator of family history of cancer (0 = No, 1 = Yes). |
| Diet_Red_Meat | Numeric index indicating the intake of red meat (0–10 scale). |
| Diet_Salted_Processed | Numeric index indicating the intake of salted and processed foods (0–10 scale). |
| Fruit_Veg_Intake | Numeric index indicating the intake of fruits and vegetables (0–10 scale). |
| Physical_Activity | Numeric index indicating overall physical activity levels (0–10 scale). |
| Air_Pollution | Numeric index representing exposure to air pollution (0–10 scale). |
| Occupational_Hazards | Numeric index indicating exposure to occupational hazards (0–10 scale). |
| BRCA_Mutation | Binary indicator of the presence of BRCA mutation (0 = No, 1 = Yes). |
| H_Pylori_Infection | Binary indicator of the presence of H. Pylori infection (0 = No, 1 = Yes). |
| Calcium_Intake | Numeric index indicating levels of calcium intake (0–10 scale). |
| Overall_Risk_Score | A composite risk metric ranging from 0 to 1, representing the overall cancer risk based on various factors. |
| Risk_Level | Categorical risk classification based on the Overall_Risk_Score: Low, Medium, or High. |
| BMI | Body Mass Index (weight in kg/height in m<sup>2</sup>), indicating whether an individual is underweight, normal weight, overweight, or obese. |

### Data preprocessing

To prepare the cancer risk factors dataset for efficient analysis and modeling, Weka data preprocessing entails a number of crucial processes. The dataset is initially loaded into Weka, which generates an ARFF representation from the CSV format. Following that, the data is checked for missing values, and affected instances are either eliminated or replicated. Distance-based techniques’ performance is then increased by scaling numerical attributes using normalization or standards. To ensure balanced training datasets, approaches for addressing class imbalance, such as SMOTE (Synthetic Minority Over-sampling Technique), are used. Following that, the data is separated into training and testing subsets so that the model can be objectively assessed [19, 20].

### Machine learning approaches

ML is important in AI because it allows computers to learn from data and generate predictions without the need for explicit programming [21]. This study investigates multiple ML techniques, with a focus on supervised learning, in which models are trained on labeled datasets containing critical features such as lifestyle choices, environmental factors, and genetic information to assess cancer risk. Supervised learning improves accuracy by identifying key risk factors and enabling comprehension of how these variables influence cancer outcomes. Furthermore, unsupervised learning approaches reveal hidden patterns in data, allowing for the development of unique patient profiles based on risk correlations. The use of advanced computer models captures complicated, nonlinear interactions between features, improving risk prediction abilities. Together, these techniques provide a solid foundation for assessing cancer risk variables, enhancing risk classification, and guiding targeted therapies to improve patient outcomes in cancer treatment [22, 23].

### Multi-layer perceptron

An input layer, one or more hidden layers, and an output layer make up the MLP, a DL neural network in which every neuron in one layer is linked to every neuron in the layer behind [24, 25]. MLPs are particularly effective at solving non-linear problems because they can learn intricate data patterns by introducing non-linearity using activation functions like sigmoid or ReLU. They function as feedforward networks, which means that information moves from input to output in a single direction. Backpropagation is used in the training process to improve accuracy by modifying weights in response to output errors. For tasks like binary or multiclass classification, the output layer produces the final predictions after each neuron acquires inputs using weighted sums and activation functions [26–28]. Fig 4 depicts the MLP diagram.

**Fig 4.**
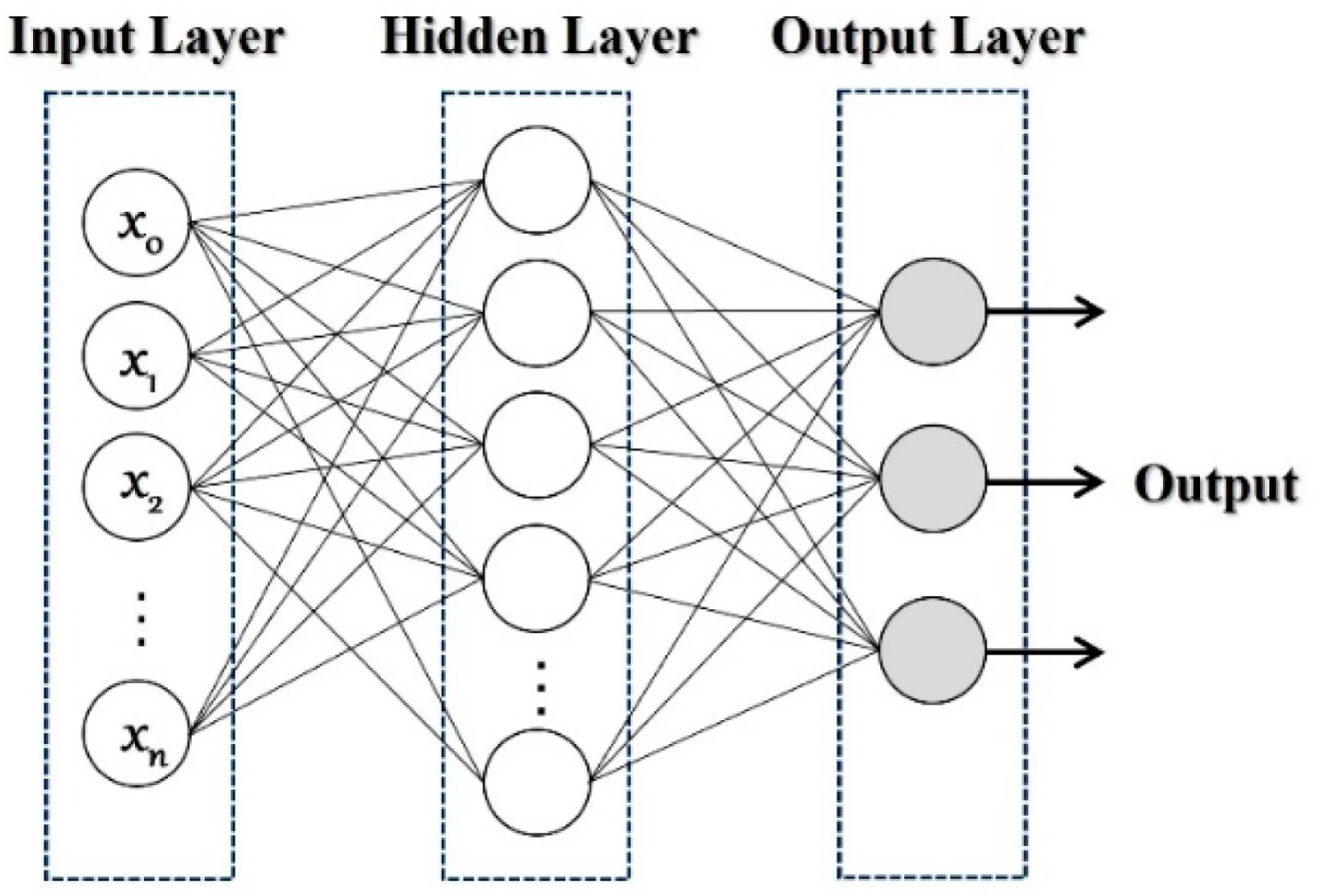
MLP structure [29].

### Random forest (RF)

A reliable ensemble learning technique that works well for both classification and regression problems is RF [30]. Using bootstrap aggregating (bagging) for data sampling, it creates several DTs during training, which avoids overfitting and improves model generalization. At each split, each tree chooses features at random, increasing diversity and lowering tree correlation as shown in Fig 5. RF averages the results for regression and employs a voting mechanism for predictions in classification. High accuracy, handling of missing values, and feature relevance scores are important benefits [31–33]. However, the towing rule, deviation, and the Gini index are important techniques for binary data division in classification issues; the Gini index is the most widely employed to quantify node impurity as shown in Eq (1).

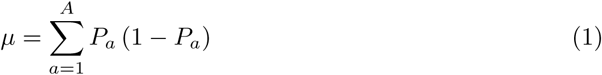

**Fig 5.**
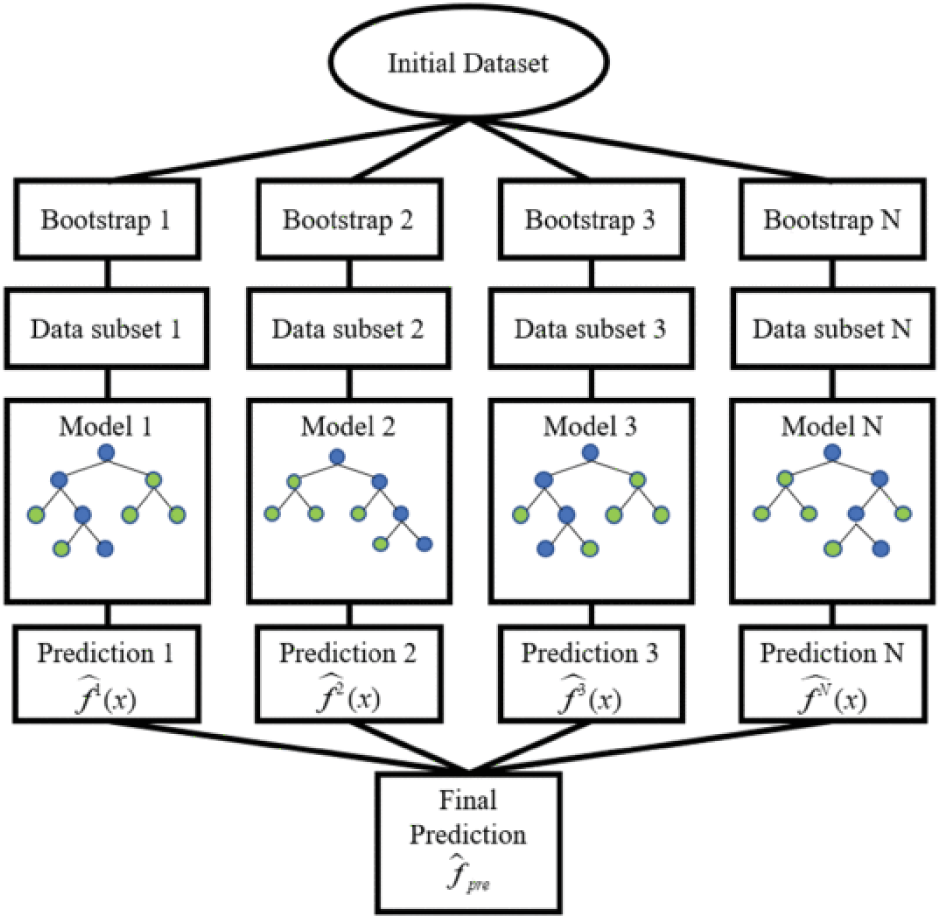
RF flow chart [36].

where the sample fraction of target class *A* is represented by *P_a_*, and a pure node made up mainly of single-class observations produces a desired *µ* value [34, 35].

### Decision tree (DT)

A supervised learning technique for regression issues is the DT algorithm. It uses decision rule learning to develop a model that predicts the value of the target variable.

The leaves of a DT indicate items which belong to the same classification, while each node chooses object attributes and specifies alternative values. The target variable separates the two forms of variable DTs: categorical and continuous [37, 38]. Based on class distributions within a dataset, the Gini impurity index *G*(*D*) measures the probability of misclassification when a class is randomly assigned. It is computed as follows for dataset *D* shown in Eq (2).

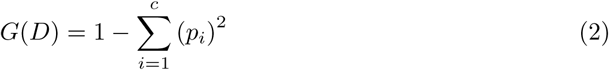

where *p_i_* represents the percentage of samples in dataset *D* that belong to class *i*, and *c* is the total number of classes [39]. A Gini impurity value of 1 denotes considerable impurity and poor classification, whereas a value of 0 implies perfect classification (all samples belong to one class).

### Logistic regression (LR)

A statistical technique for binary classification called LR predicts the possibility that an input will be classified into a particular category. It transforms linear combinations of input information into probabilities between 0 and 1 using the logistic function. The key is maximum likelihood estimation (MLE), which maximizes the likelihood of observed data to optimize model parameters [40, 41].

In logistic regression, the odd—that is, the probability of success divided by the probability of failure—is translated using the logit formula. Eq (3) and Eq (4) depict this logistic function, which is also referred to as the log odds or the natural logarithm of odds:

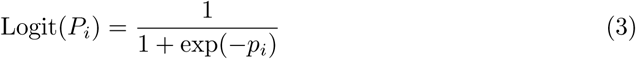

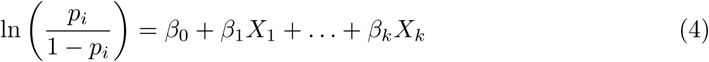

The independent variable in this LR equation is *x*, while the dependent or response variable is logit(*p_i_*). Maximum likelihood estimation (MLE) is frequently used to estimate the beta parameter or coefficient in this model [42].

### Naive Bayes (NB)

Based on Bayes’ theorem, NB is a probabilistic classifier that works well for binary and multi-class classification, especially in high-dimensional fields. Despite possible violations of this condition, its “naive” assumption of independent features simplifies modeling and frequently yields robust results. The classifier chooses the class with the highest probability after assessing the probabilities of each class and feature.

Computational efficiency, scalability, and interpretability are advantages; correlated feature errors and difficulties with unbalanced datasets are disadvantages. In general, NB is a simple and popular classification method [43, 44]. Eq (5) illustrates how the Bayesian classification framework determines the posterior probability.

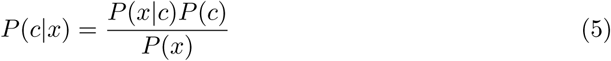

where *P* (*x*) is the evidence, *P* (*x|c*) is the likelihood distribution, *P* (*c|x*) is the posterior probability, *x* is the feature vector, and *c* is the classification variable [45].

## Performance evaluation

### Confusion matrix

A classification model’s efficacy can be assessed using a confusion matrix, a performance measurement tool. It offers a thorough comparison of the actual and predicted classes, making it easy to assess the model’s performance [46]. However, the confusion matrix consists of four parts as shown in Table 3:

- True Positives (TP): The number of cases that the model properly predicts as positive [47].
- True Negatives (TN): The number of cases that the model properly predicts as negative [48].
- False Positives (FP): The quantity of cases that the model mistakenly predicts as positive [49].
- False Negatives (FN): The quantity of cases that are mistakenly projected to be negative because the model is unable to recognize a positive class [50].

**Table 3.** Confusion matrix [51].

| Actual | Predicted |  |
| --- | --- | --- |
|  | True Positive (TP) | False Negative (FN) |
|  | False Positive (FP) | True Negative (TN) |

In this work a multi-class confusion matrix is used for assessing the effectiveness of classification models, particularly when dealing with three or more classes. It provides a detailed summary of how the model’s predictions compare to the actual classifications in the dataset. In the context of cancer risk assessment, where categories could include High, Medium, and Low risk, the multi-class confusion matrix provides insights about the model’s advantages and disadvantages in distinguishing between various levels.

Fig 6(a) depicts the NB confusion matrix, Fig 6(b) the RF confusion matrix, Fig 6(c) the DT confusion matrix, Fig 6(d) the MLP confusion matrix, and Fig 6(e) the LR confusion matrix. The multi-class confusion matrix for three risk categories (High, Medium, and Low) is organized as demonstrated in Table 4.

**Fig 6.**
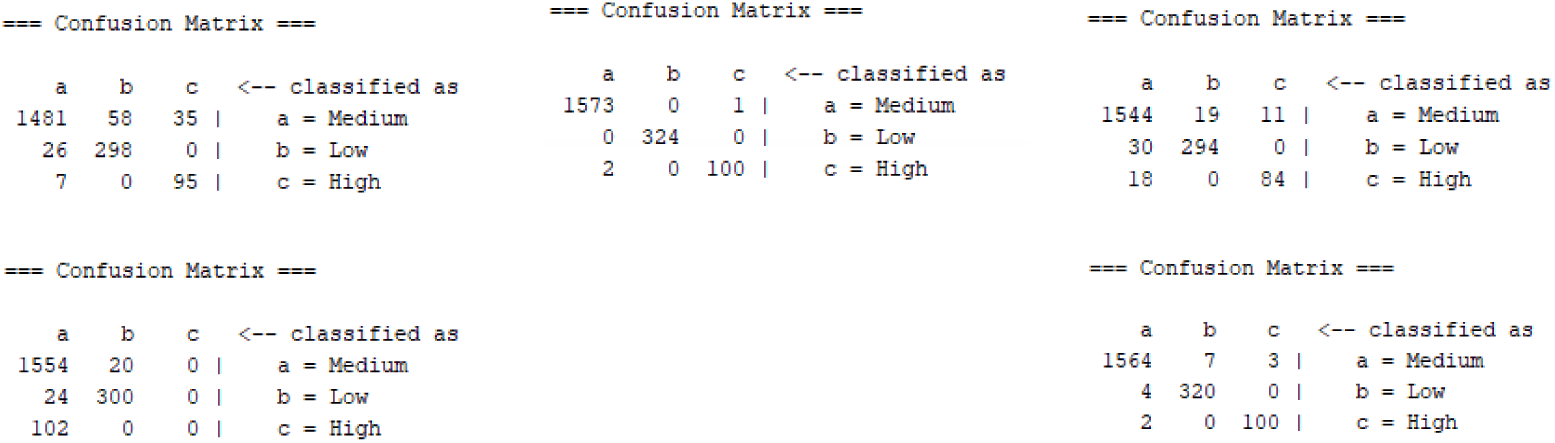
Confusion matrix of all classifiers. (a) confusion matrix of NB classifier, (b) confusion matrix of RF classifier, (c) confusion matrix of DT classifier, (d) confusion matrix of MLP classifier, and (e) confusion matrix of LR classifier.

**Table 4.** Multi-class confusion matrix for three cancer risk factors.

|  |  | Predicted |  |  |
| --- | --- | --- | --- | --- |
| Actual |  | High (H) | Medium (M) | Low (L) |
|  | High (H) | TP_H | FN_M | FN_L |
|  | Medium (M) | FP_H | TP_M | FN_L |
|  | Low (L) | FP_H | FP_M | TP_L |

### Performance matrices

In this section, the efficacy of the classification models of cancer risk variables was assessed using four critical performance metrics: accuracy, F-measure, precision, and sensitivity of performance metrics as shown in Table 5. Each indicator provides distinct insights into the models’ capabilities and predictive accuracy. However, the measurements enable well-informed comparisons between various algorithms and offer an extensive understanding of how effectively the models predicted results [52]. Among the main indicators of performance used are:

- Accuracy: The ratio of accurately predicted instances (TP + TN) to total instances indicates the model’s overall reliability [53].
- Precision: The ratio of genuine positives to all predicted positives (TP / (TP + FP)) represents the accuracy of positive predictions [54].
- Recall (Sensitivity): The ratio of true positives to total real positives (TP / (TP + FN)) indicates the model’s capacity to detect positive cases [55].
- F1-measure: 2 *×* (Precision *×* Sensitivity) / (Precision + Sensitivity) is the harmonic mean of accuracy and recall, providing a balance between the two [56].

**Table 5.**
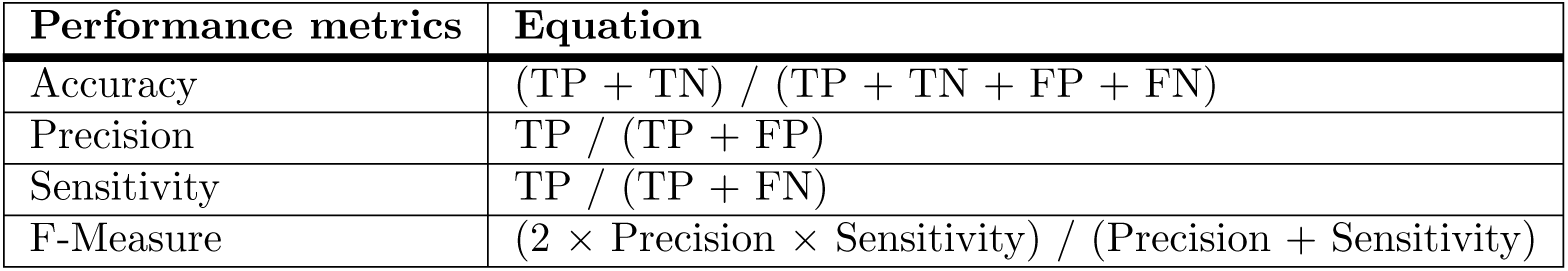
Performance metrics [57].

### Cross-validation

Cross-validation is used in statistics and ML to pick models and parameters, as well as to ensure accuracy by fitting and testing models on diverse data sets. Traditional methods, such as V-fold and leave-one-out, frequently result in overfitting [43].

Cross-validation often does not allow for a training-testing split ratio in low-dimensional linear models, which are widely utilized due to larger training samples [58]. This study uses the 10-fold cross-validation approach, in which the dataset is randomly divided into ten equal parts: nine for training and one for validation as shown in Fig 7. This technique is repeated ten times to provide an average performance metric. The 10-fold cross-validation technique entails rearranging the data and assuring randomization in the split. The model is trained on nine folds for each fold before being evaluated on the remaining fold to capture performance measures such as accuracy. This repeatable process improves the accuracy of model performance predictions, reduces overfitting, and makes the best use of limited data size [59–62].

**Fig 7.**
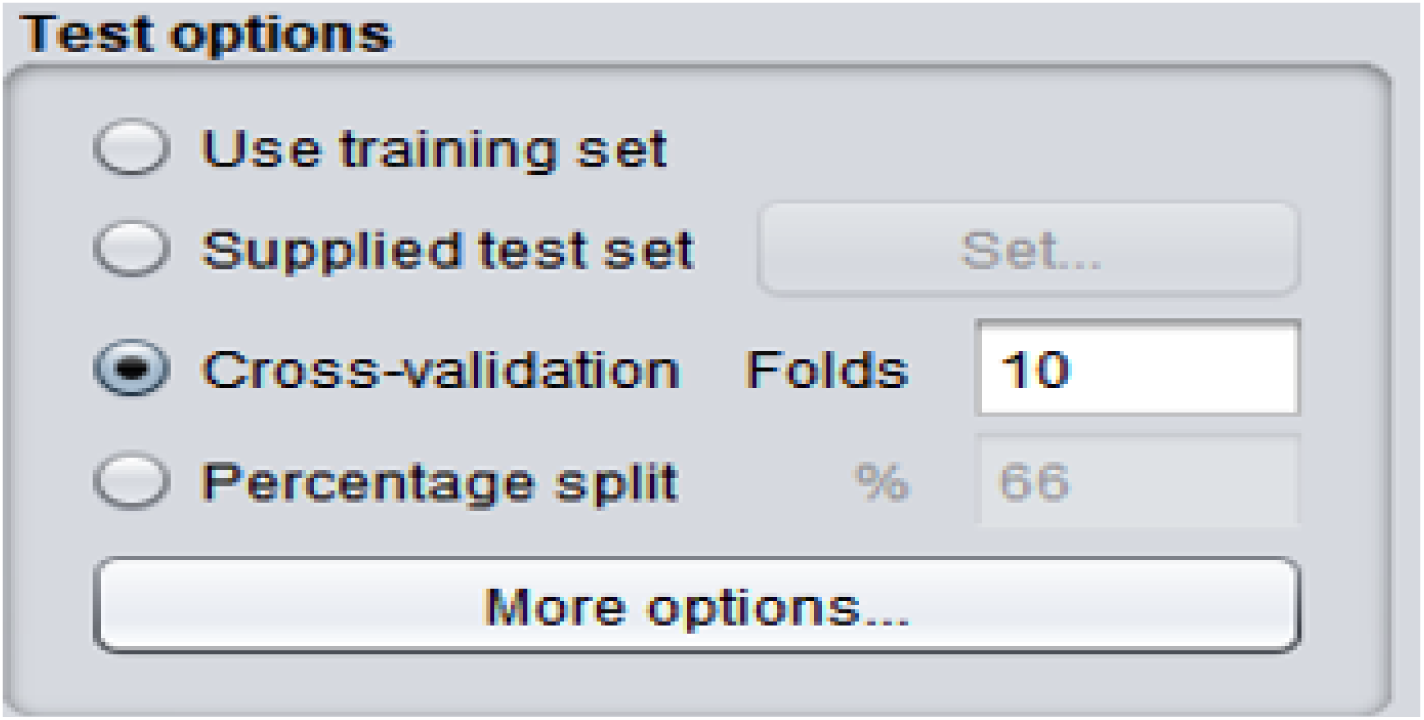
10-fold cross validation setting.

## Results and discussion

In ML, model evaluation is critical for determining the effectiveness of algorithms in predictions. This discussion compares five classification models using metrics like accuracy, F-measure, precision, and sensitivity, with each providing unique insights into model performance and guiding algorithm selection for specific tasks.

Accuracy is a crucial parameter for determining a model’s overall validity in predicting target classes. Among the models tested, the RF performs well with an accuracy of 99.85%, suggesting its great reliability and effectiveness for the provided dataset. LR and DT both perform well, with accuracies of 99.20% and 96.10%, respectively. In contrast, NB and MLP have lower accuracy rates of 93.70% and 92.70%, indicating that these models may be less stable or robust under certain data conditions. Fig 8 depicts the accuracy of all classifiers.

**Fig 8.**
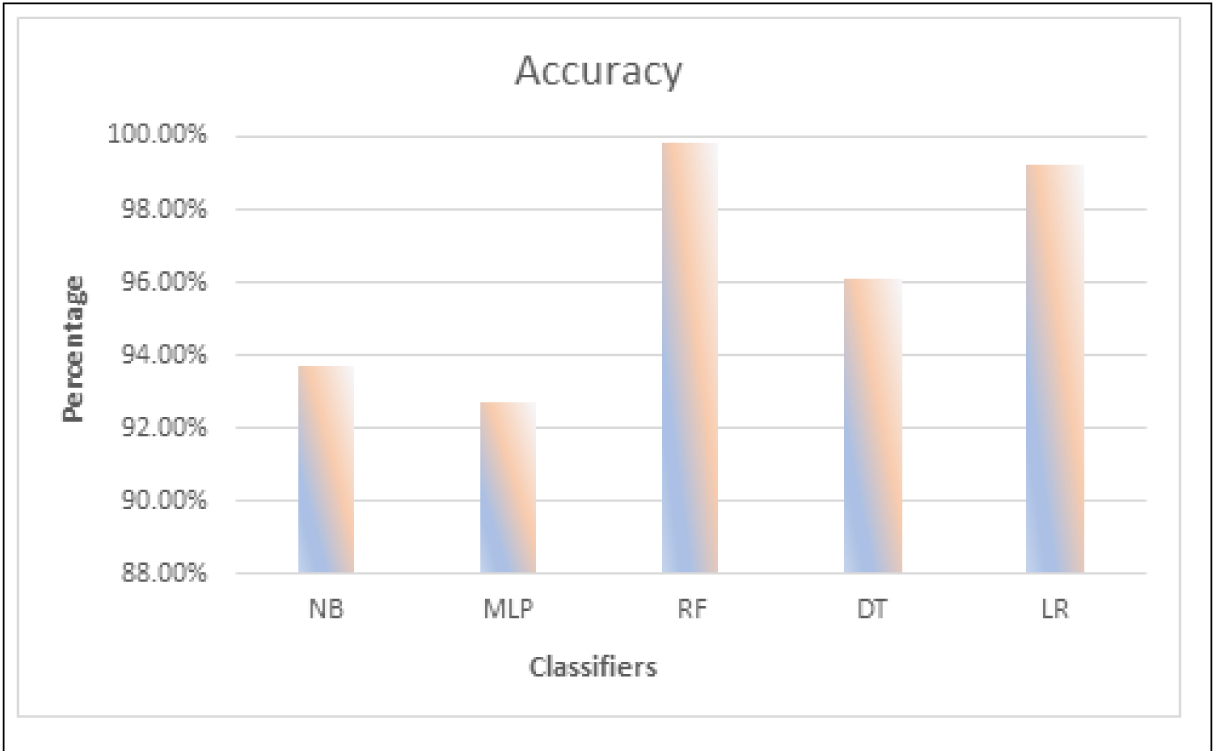
Accuracy of all classifiers.

Furthermore, the F-measure is critical for assessing the balance of precision and recall, particularly in datasets with class imbalances. The RF model performs highly once more, with an F-measure of 99.80%, indicating that it can retain both precision and recall. LR comes in second with 99.20%, followed by MLP with a commendable 96.35%. NB and DT had lower F-measures of 93.90% and 96.10%, respectively, indicating that these models may not handle the decision between true positives and false positives as efficiently. Fig 9 depicts the F-measure of all classifiers.

**Fig 9.**
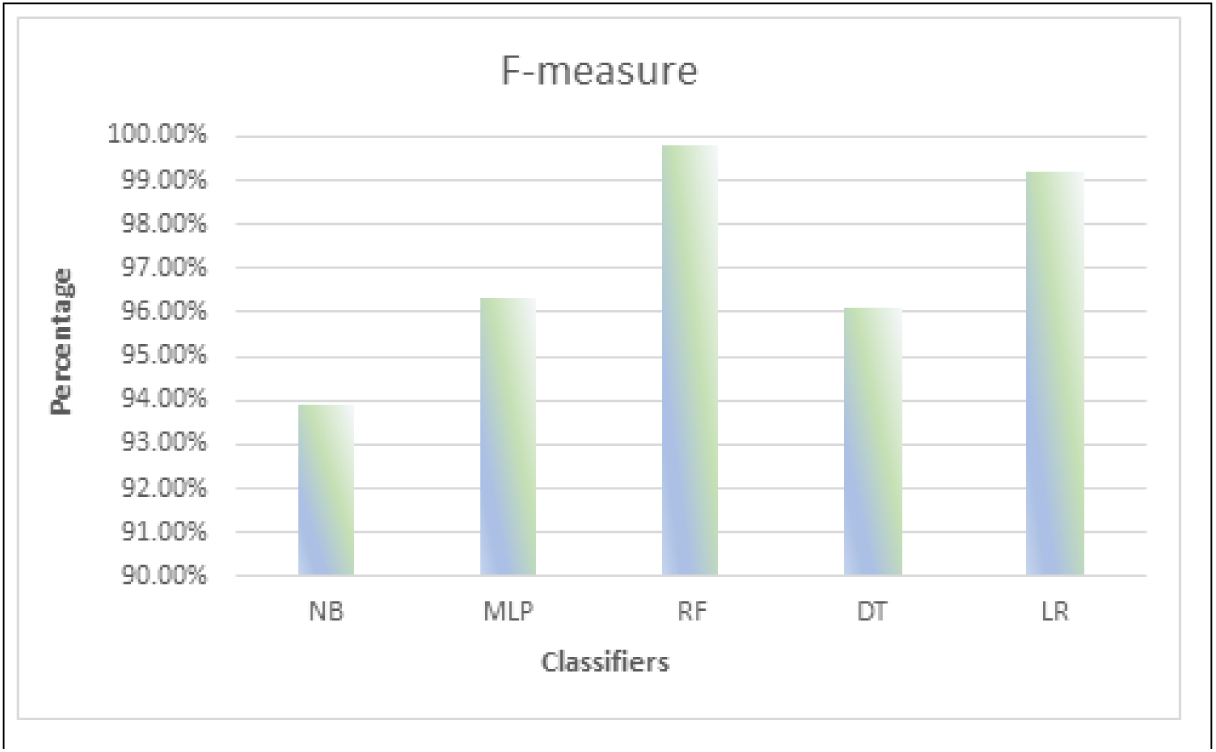
F-measure of all classifiers.

Precision represents the model’s capacity to make relevant positive predictions while limiting the amount of false positives. Both RF and LR have high precision ratings of 99.80%, demonstrating their efficacy in filtering out irrelevant cases. However, the MLP has a lower precision of 93.15%, indicating that it may misclassify more occurrences as positive than the others. NB and DT have similar precision levels of 94.30% and 96.10%, indicating that while they perform well, there is potential for advancement.

Fig 10 depicts the precision of all classifiers.

**Fig 10.**
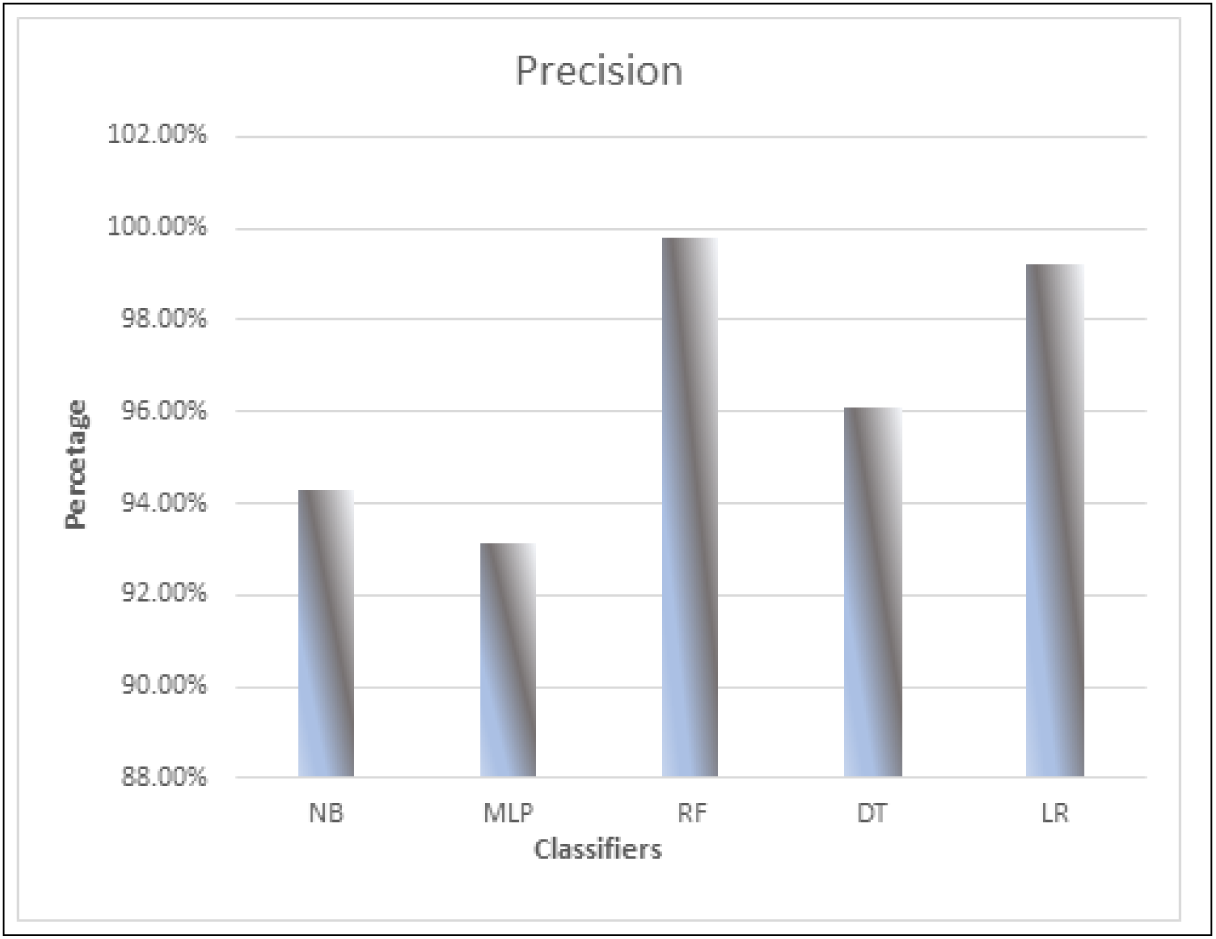
Precision of all classifiers.

Finally, sensitivity (or recall) assesses the model’s ability to correctly identify genuine positive cases. In this aspect, RF leads with a sensitivity of 99.90%, closely followed by LR at 99.20%, demonstrating their ability to detect true positives successfully. DT came in second with 96.10%, while MLP and NB trail with sensitivity of 92.70% and 93.70%, respectively. This variation implies that RF and LR perform much better at limiting missed opportunities to find positive instances, which is crucial in applications where positive examples are critical. Fig 11 depicts the sensitivity of all classifiers.

**Fig 11.**
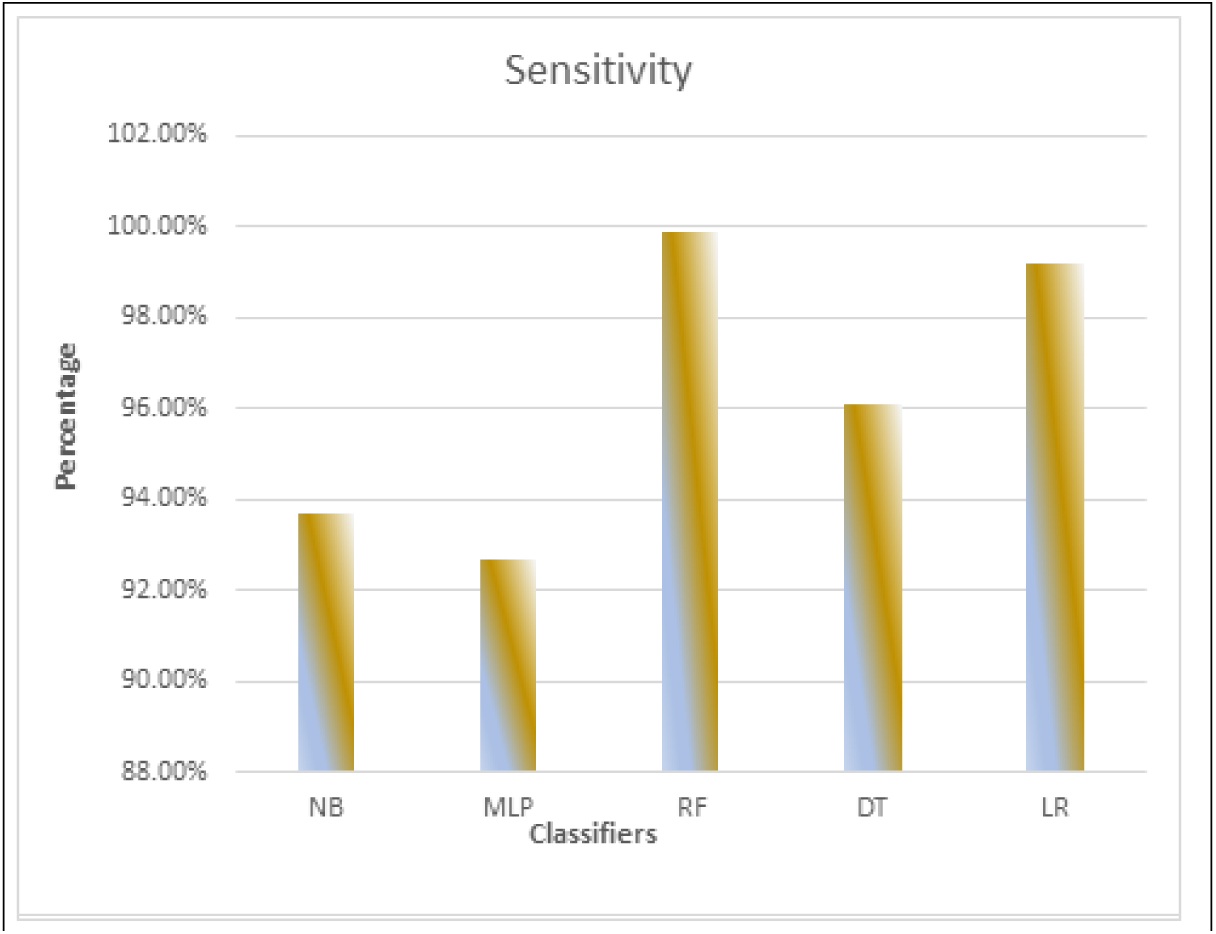
Sensitivity of all classifiers.

Based on the performance metrics tested, the RF model outperforms the other classifiers. With high scores across all parameters, it exhibits an exceptional capacity to reliably detect both positive and negative cases while minimizing misclassifications. Its constant performance demonstrates its dependability for a wide range of classification tasks, making it an excellent choice for applications where accuracy and robustness are critical. This analysis emphasizes the significance of using RF as a preferable approach for scenarios that require adequate classification outcomes. Table 6 presents the performance evaluation comparison of all classifiers while Fig 12 presents the comparative analysis of different classifiers’ performances.

**Fig 12.**
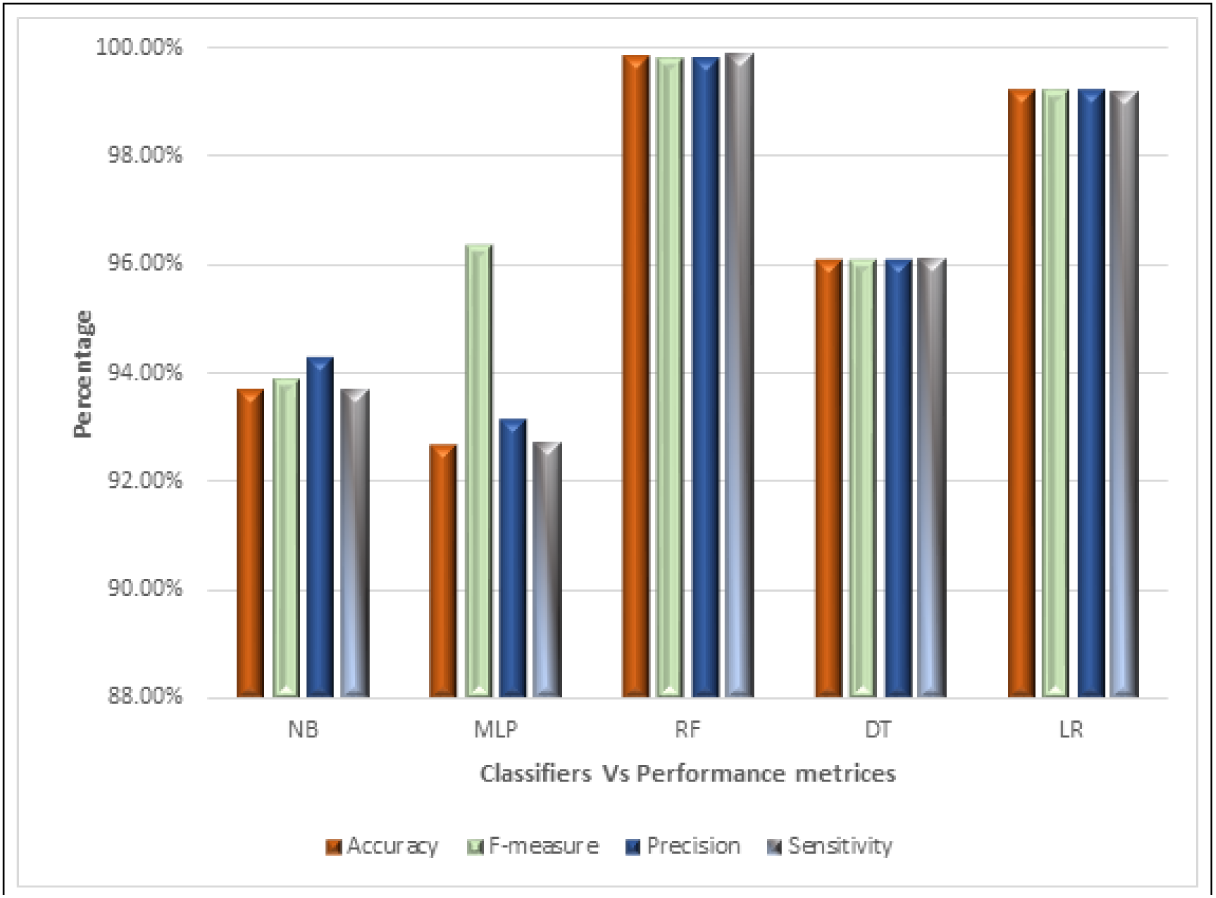
Comparative analysis of different classifiers’ performances.

**Table 6.** Performance evaluation comparison of all classifiers.

| Model | Accuracy | F-measure | Precision | Sensitivity |
| --- | --- | --- | --- | --- |
| NB | 93.70% | 93.90% | 94.30% | 93.70% |
| MLP | 92.70% | 96.35% | 93.15% | 92.70% |
| RF | 99.85% | 99.80% | 99.80% | 99.90% |
| DT | 96.10% | 96.10% | 96.10% | 96.10% |
| LR | 99.20% | 99.20% | 99.20% | 99.20% |

## Conclusion

Insights on key cancer risk factors may support personalized healthcare interventions. This work highlights the revolutionary impact of AI in cancer diagnosis, urging further research to increase model applicability across various populations and test novel algorithms for improved prediction accuracy. Using a vast Kaggle dataset, the study demonstrates the significant effect of AI and ML on cancer risk assessment, supporting successful healthcare practices. The study used a Java-based ML framework to develop and analyze multiple prediction models, with a particular emphasis on the RF method because of its ability to handle complicated datasets and interpretability, which is critical in clinical settings. The RF model attained an excellent accuracy of 99.85% using 10-fold cross-validation, proving its suitability for real-world applications. The results support a data-driven approach to early cancer identification, allowing healthcare providers to implement individualized care strategies that improve patient outcomes. AI-powered solutions have the ability to go beyond prediction by encouraging proactive steps that respond to individual risk variables. Cancer care can be made more personalized and focused by utilizing such technology, thereby boosting treatment quality and intervention effectiveness. Future research in predictive modeling is required to fit developments with changing patient care requirements.

## Future work

Future AI and ML research for cancer risk assessment should focus on a number of critical areas in order to improve prediction models. Key objectives include extending datasets to represent varied populations, which improves generalizability across demographics such as race, age, and socioeconomic level. Advanced techniques such as DL and ensemble approaches are encouraged to improve predicting accuracy by recognizing complex patterns in data. Integrating longitudinal data is critical for dynamic risk assessments that respond to patients’ changing health profiles. Improved feature engineering led by domain experience can help to tweak model performance further. It is essential that healthcare professionals participate in the practical application of these models to guarantee compliance with existing procedures and compliance to ethical principles. In addition, developing systems that provide individualized preventative and early treatment recommendations will improve patient care. Finally, long-term evaluations of AI-driven risk assessments on patient outcomes are essential for validating their effectiveness over time. Addressing these issues will considerably improve the use of AI and ML in cancer risk assessment, ultimately leading to better patient care in oncology.

## Data availability

The dataset utilized in this study, “Cancer Risk Factors Dataset,” authored by T. Masry and published in 2023, is available at https://www.kaggle.com/datasets/tarekmasryo/cancer-risk-factors-dataset. This resource was accessed on October 17, 2023, and can be cited as: T. Masry, “Cancer Risk Factors Dataset,” Kaggle, 2023.

## Acknowledgments

The authors want to acknowledge the financial support under the grant from Multimedia University (MMU) Postdoc Fellowship Fund under the grant number MMUI/260011.

## Author Contribution

Areen Arabiat contributed to the conceptualization, methodology, software development, validation, formal analysis, investigation, resource management, data curation, original draft writing, review and editing, visualization, supervision, and funding acquisition of the manuscript. Hamza Abu Owida was involved in the original draft writing, conceptualization, validation, formal analysis, investigation, resource management, review and editing, visualization and project administration. The authors state no funding involved and no conflict of interest.

## References

1. Sung H, Ferlay J, Siegel RL, Laversanne M, Soerjomataram I, Jemal A, et al. Global Cancer Statistics 2020: GLOBOCAN estimates of incidence and mortality worldwide for 36 cancers in 185 countries. CA Cancer J Clin. 2021;71(3):209–249.

2. Vineis P, Wild CP. Global cancer patterns: causes and prevention. Lancet. 2014;383(9916):549–557.

3. Anand P, Kunnumakkara AB, Sundaram C, Harikumar KB, Tharakan ST, Lai OS, et al. Cancer is a preventable disease that requires major lifestyle changes. Pharm Res. 2008;25(9):2097–2116.

4. Song M, Giovannucci EL. Cancer risk: many factors contribute. Science. 2015;347(6223):728–729.

5. Hippisley-Cox J, Coupland C. Development and validation of risk prediction algorithms to estimate future risk of common cancers in men and women: prospective cohort study. BMJ Open. 2015;5(3):e007825.

6. Libbrecht MW, Noble WS. Machine learning applications in genetics and genomics. Nat Rev Genet. 2015.

7. Kourou K, Exarchos TP, Exarchos KP, Karamouzis MV, Fotiadis DI. Machine learning applications in cancer prognosis and prediction. Comput Struct Biotechnol J. 2015.

8. Wainberg M, Merico D, Delong A, Frey BJ. Deep learning in biomedicine. Nat Biotechnol. 2018.

9. Rajkomar A, Dean J, Kohane I. Machine learning in medicine. N Engl J Med. 2019.

10. Kinar Y, Kalkstein N, Akiva P, Levin B, Half EE, Goldshtein I, et al. Development and validation of a predictive model for detection of colorectal cancer in primary care by analysis of complete blood counts: a binational retrospective study. J Am Med Inform Assoc. 2016;23(5):879–890.

11. Kinar Y, Akiva P, Choman E, Kariv R, Shalev V, Levin B, et al. Performance analysis of a machine learning flagging system used to identify a group of individuals at a high risk for colorectal cancer. PLoS One. 2017;12(2):e0171759.

12. Hilsden RJ, Heitman SJ, Mizrahi B, Narod SA, Goshen R. Prediction of findings at screening colonoscopy using a machine learning algorithm based on complete blood counts (ColonFlag). PLoS One. 2018;13(11):e0207848.

13. Ke TM, Lophatananon A, Muir KR. An integrative pancreatic cancer risk prediction model in the UK Biobank. Biomedicines. 2023;11(12):3206.

14. Tu H, Zhao Y, Cui J, Lu W, Sun G, Xu X, et al. Improving lung cancer risk prediction using machine learning: a comparative analysis of stacking models and traditional approaches. Cancers (Basel). 2025;17(10):1651.

15. Chiu PKF, Shen X, Wang G, Ho CL, Leung CH, Ng CF, et al. Enhancement of prostate cancer diagnosis by machine learning techniques: an algorithm development and validation study. Prostate Cancer Prostatic Dis. 2022;25:672–676.

16. Dianati-Nasab M, Salimifard K, Mohammadi R, Saadatmand S, Fararouei M, Hosseini KS, et al. Machine learning algorithms to uncover risk factors of breast cancer: insights from a large case-control study. Front Oncol. 2023;13:1276232.

17. Guo Q, Wu P, He J, Zhang G, Zhou W, Chen Q, et al. Machine learning algorithms predict breast cancer incidence risk: a data-driven retrospective study based on biochemical biomarkers. BMC Cancer. 2025;25:1061.

18. Masry T. Cancer Risk Factors Dataset. Kaggle; 2023. Available from: https://www.kaggle.com/datasets/tarekmasryo/cancer-risk-factors-dataset.

19. Awojoyogbe BO, Dada MO. Performance Evaluation of Machine Learning Classification of Brain Tumors with WEKA and Python Programming. In: Series in Bioengineering. 2024. p. 229–246. doi:10.1007/978-981-97-6370-29.

20. Arabiat A, Hassan M, Almomani O. WEKA-based machine learning for traffic congestion prediction in Amman City. IAES Int J Artif Intell. 2024;13(4):4422–4434. doi:10.11591/ijai.v13.i4.pp4422-4434.

21. Almomani O, et al. A Robust Model for Android Malware Detection via ML and DL classifiers. Mesopotamian J Big Data. 2025;2025:261–277. doi:10.58496/mjbd/2025/017.

22. Rolf B, et al. A review on unsupervised learning algorithms and applications in supply chain management. Int J Prod Res. 2024;63(5):1933–1983. doi:10.1080/00207543.2024.2390968.

23. Zhang J, Yang L, Mohammadabadi SMS, Yan F. A survey on self-supervised learning: Recent advances and open problems. Neurocomputing. 2025;655:131409. doi:10.1016/j.neucom.2025.131409.

24. Setitra MA, Fan M, Agbley BLY, Bensalem ZEA. Optimized MLP-CNN Model to Enhance Detecting DDoS Attacks in SDN Environment. Network. 2023;3(4):538–562. doi:10.3390/network3040024.

25. Arabiat A, Hassan M, Al Momani O. Traffic congestion prediction using machine learning: Amman City case study. In: MIEITS 2024. vol. 13188. SPIE; 2024. p. 38–45. doi:10.1117/12.3030849.

26. Ahmed S. A Software Framework for Predicting the Maize Yield Using Modified Multi-Layer Perceptron. Sustainability. 2023;15(4):3017. doi:10.3390/su15043017.

27. Hammad MM. Deep Learning activation functions: Fixed-Shape, Parametric, Adaptive, Stochastic, Miscellaneous, Non-Standard, Ensemble. arXiv; 2024. doi:10.48550/arxiv.2407.11090.

28. Mohammadzadeh A, Sabzalian MH, Castillo O, Sakthivel R, El-Sousy FFM, Mobayen S. Multilayer Perceptron (MLP) Neural Networks. In: Neural Networks and Learning Algorithms in MATLAB. Springer, Cham; 2022. doi:10.1007/978-3-031-14571-12.

29. Lee H, Kim D, Gu JH. Prediction of food factory energy consumption using MLP and SVR algorithms. Energies. 2023;16(3):1550. doi:10.3390/en16031550.

30. Lachaud A, Adam M, Mǐskovíc I. Comparative Study of Random Forest and Support Vector Machine Algorithms in Mineral Prospectivity Mapping with Limited Training Data. Minerals. 2023;13(8):1073. doi:10.3390/min13081073.

31. Wang J, Ma S, Jiao P, Ji L, Sun X, Lu H. Analyzing the Risk Factors of Traffic Accident Severity Using a Combination of Random Forest and Association Rules. Appl Sci. 2023;13(14):8559. doi:10.3390/app13148559.

32. Arabiat A, Altayeb M, Alkhrissat T. Innovative AI Approaches for concrete Strength Prediction: Towards Sustainable Buildings. Civ Eng Archit. 2025;13(6):4254–4265. doi:10.13189/cea.2025.130612.

33. Nikhitha KV, Bhavya K, Nandini DU. Fake Account Detection on Social Media using Random Forest Classifier. In: ICICCS 2023. 2023. p. 806–811. doi:10.1109/ICICCS56967.2023.10142841.

34. Chen C, Liang J, Sun W, Yang G, Meng X. An automatically recursive feature elimination method based on threshold decision in random forest classification. Geo-spatial Inf Sci. 2024;28(4):1494–1519. doi:10.1080/10095020.2024.2387457.

35. Bafail O. Optimizing Smart City Strategies: A Data-Driven Analysis using random forest and regression analysis. Appl Sci. 2024;14(23):11022. doi:10.3390/app142311022.

36. Kwon MS, Lim DK. A study on the optimal design of PMA-SYNRM for electric vehicles combining random forest and genetic algorithm. IEEE Access. 2023;11:52357–52369. doi:10.1109/access.2023.3279126.

37. Rahman MSA, Jamaludin NAA, Zainol Z, Sembok TMT. The application of Decision Tree Classification Algorithm on Decision-Making for Upstream Business. Int J Adv Comput Sci Appl. 2023;14(8). doi:10.14569/ijacsa.2023.0140873.

38. Mienye ID, Jere N. A Survey of Decision Trees: Concepts, Algorithms, and Applications. IEEE Access. 2024;12:86716–86727. doi:10.1109/ACCESS.2024.3416838.

39. Nguyen TD, et al. Efficient and Explainable Bearing Condition Monitoring with Decision Tree-Based Feature Learning. Machines. 2025;13(6):467. doi:10.3390/machines13060467.

40. Zhang H, Su L, Liu Y, et al. Assessment of the effects of characterization methods selection on the landslide susceptibility: a comparison between logistic regression (LR), naive bayes (NB) and radial basis function network (RBF Network). Bull Eng Geol Environ. 2025;84:134. doi:10.1007/s10064-025-04097-2.

41. Hua Y, Stead TS, George A, Ganti L. Clinical Risk Prediction with Logistic Regression: Best Practices, Validation Techniques, and Applications in Medical Research. Acad Med Surg. 2025. doi:10.62186/001c.131964.

42. Arabiat A, Altayeb M. An automated system for classifying types of cerebral hemorrhage based on image processing techniques. Int J Electr Comput Eng. 2024;14(2):1594–1603. doi:10.11591/ijece.v14i2.pp1594-1603.

43. Hassan M, Arabiat A. An evaluation of multiple classifiers for traffic congestion prediction in Jordan. Indones J Electr Eng Comput Sci. 2024;36(1):461–468. doi:10.11591/ijeecs.v36.i1.pp461-468.

44. Alshammari AH, Bencsik G, Ali AH. A survey of six classical classifiers, including algorithms, methodological characteristics, foundational variants, and recent advances. Algorithms. 2026;19(1):37. doi:10.3390/a19010037.

45. Kaur S, et al. High-accuracy lung disease classification via logistic regression and advanced feature extraction techniques. Egypt Inform J. 2024;29:100596. doi:10.1016/j.eij.2024.100596.

46. Zeng G. Invariance Properties and Evaluation Metrics Derived from the Confusion Matrix in Multiclass Classification. Mathematics. 2025;13(16):2609. doi:10.3390/math13162609.

47. Gautam D, Mawardi Z, Elliott L, Loewensteiner D, Whiteside T, Brooks S. Detection of invasive species (Siam weed) using Drone-Based Imaging and YOLO Deep Learning model. Remote Sens. 2025;17(1):120. doi:10.3390/rs17010120.

48. Rosas-Diaz M, et al. New tool against tuberculosis: The potential of the LAMP Lateral Flow assay in Resource-Limited Settings. Curr Issues Mol Biol. 2025;47(8):585. doi:10.3390/cimb47080585.

49. Zhu L, et al. Improving the precision of Deep-Learning-Based head and neck target Auto-Segmentation by leveraging radiology reports using a large language model. Cancers. 2025;17(12):1935. doi:10.3390/cancers17121935.

50. Kovalchuk AV, Lebedev AA, Shemagina OV, Nuidel IV, Yakhno VG, Stasenko SV. Enhancing cascade object detection accuracy using correctors based on High-Dimensional Feature Separation. Technologies. 2025;13(12):593. doi:10.3390/technologies13120593.

51. Arabiat AM, Eljaafreh YG. Intrusion Detection in Wireless Sensor Networks Using ML Based Classification of Denial of Service (DoS) Attacks. J Commun. 2025;20(4).

52. Abualhaj M. Spam detection boosted by Firefly-Based feature selection and optimized classifiers. Int J Adv Soft Comput Appl. 2025;17(3). doi:10.15849/ijasca.251130.02.

53. Owida HA, Arabiat A, Al-Ayyad M, Altayeb M. Advancements in machine learning techniques for precise detection and classification of lung cancer. Bull Electr Eng Inform. 2025;14(6):4521–4533. doi:10.11591/eei.v14i6.10527.

54. Crossa J, et al. Machine learning algorithms translate big data into predictive breeding accuracy. Trends Plant Sci. 2024;30(2):167–184. doi:10.1016/j.tplants.2024.09.011.

55. Bilal M, Shah AA, Abbas S, Khan MA. High-Performance deep learning for instant pest and disease detection in precision agriculture. Food Sci Nutr. 2025;13(9):e70963. doi:10.1002/fsn3.70963.

56. Hosseinzadeh M, et al. Improving phishing email detection performance through deep learning with adaptive optimization. Sci Rep. 2025;15(1):36724. doi:10.1038/s41598-025-20668-5.

57. Abdu A, et al. Cross-project software defect prediction based on the reduction and hybridization of software metrics. Alex Eng J. 2024;112:161–176. doi:10.1016/j.aej.2024.10.034.

58. Owida HA, et al. A deep learning-based dual-branch framework for automated skin lesion segmentation and classification via dermoscopic Images. Sci Rep. 2025;15(1):37823. doi:10.1038/s41598-025-21783-z.

59. Arabiat AM. Intelligent model for detecting GAN-Generated images based on Multi-Classifier and advanced data mining techniques. Int J Electr Electron Eng Telecommun. 2024;14(3):147–157. doi:10.18178/ijeetc.14.3.147-157.

60. Jin B, Xu X. Predicting wholesale edible oil prices through Gaussian process regressions tuned with Bayesian optimization and cross-validation. Asian J Econ Bank. 2024;9(1):64–82. doi:10.1108/ajeb-06-2024-0070.

61. Seraj A, et al. Cross-validation. In: Elsevier eBooks. 2023. p. 89–105. doi:10.1016/b978-0-12-821285-1.00021-x.

62. Abu-Shareha AA, et al. A comparative study of the diabetes progression prediction techniques. Discov Artif Intell. 2025. doi:10.1007/s44163-025-00770-3.

